# Comparative efficacy and safety of anti-cryptosporidial agents: An *in vitro* study on Nitazoxanide, HFL, KDU731, and Paromomycin against *Cryptosporidium parvum*

**DOI:** 10.1101/2024.02.09.579383

**Authors:** Saffron T.G. Whitta, Bridget Lamont, Rossarin Suwanarusk, Bruce M. Russell, Morad-Rémy Muhsin-Sharafaldine

## Abstract

This study evaluates the *in vitro* effectiveness of the anti-cryptosporidial agents Nitazoxanide, Halofuginone, the pyrazolopyridine analogue KDU731, and Paromomycin in combating the significant zoonotic pathogen *Cryptosporidium parvum*. The study utilizes HCT-8 host cells to culture *C. parvum* and fluorescent microscopy and qPCR for detecting parasitic growth. The efficacy of the compounds was assessed by calculating their inhibitory concentrations against the total growth of *C. parvum* at 48 hours post-infection. The study further investigates the impact of these compounds on early parasitophorous vacuole formation, merozoite egress, host cell viability, and cell growth cycle. KDU731 displayed the most promising profile, with low nanomolar (102 nM ± 2.28) activity and negligible host cell toxicity. This study offers new insights into the relative efficacy and safety of various anti-cryptosporidial compounds, highlighting their stage-specific effects on *C. parvum* and the consequential impacts on host cells. Identifying safe and effective anti-cryptosporidial agents contributes significantly to the One Health approach, emphasizing the importance of integrated strategies in controlling zoonotic diseases.

## Introduction

Cryptosporidiosis is a parasitic disease of the small intestine and respiratory tract, caused by the protozoan parasite *Cryptosporidium* spp., affecting human populations and domestic and wild animals.^1^ Globally, approximately 84% of all deaths caused by cryptosporidiosis are of children under the age of five.^2^ Children in developing countries show greater risk of severe symptoms such as vomiting, dehydration, fever, prolonged diarrhea and even death due to *Cryptosporidium* infection.^2,3^ Moreover, as severe symptoms are prolonged in impoverished children, long lasting health issues can occur, such as stunted growth and cognitive issues.^4,5^ More, *Cryptosporidium*-infected children under one year old are unable to catch up with lost growth, especially due to the severe malnutrition/infection cycle exacerbating the developmental delay.^6,7^ Of all species that cause cryptosporidiosis in humans, *C. parvum* is of interest due to its high incidence rate and zoonotic transmission between livestock (beef and lamb) and humans.^8^

Unfortunately, there is neither vaccine nor effective treatment to treat cryptosporidiosis. The antimicrobial thiazolide, nitazoxanide, is the only approved treatment against cryptosporidiosis in humans.^9,10^ Nitazoxanide was initially discovered as a drug to treat tapeworm infection, later being repurposed as a drug to treat both *Cryptosporidium spp*. and *Giardia intestinalis* infections.^11,12^ Studies have suggested that nitazoxanide works by disrupting the parasite’s anaerobic respiration by blocking the pyruvate/ferredoxin oxidoreductase enzyme-dependent electron transfer reaction.^12^ While Nitazoxanide is moderately efficacious in immunocompetent patients, its efficacy is highly reduced in immunocompromised patients, children, and absent in those with HIV.^10^

Animal cryptosporidiosis also poses a large threat, primarily to livestock farming with estimated millions in losses due to infection.^13-15^ Currently the only drug for treatment of livestock with cryptosporidiosis is the synthetic quinazolinone halofuginone lactate (HFL).^16,17^ HFL was initially shown to inhibit cancerous tumors and was pushed as a cancer drug, only to be later repurposed as an antiprotozoal drug for treatment of *Cryptosporidiosis* in livestock.^18^ It has been found that as a prophylactic HFL can both delay oocyst shedding and reduce symptoms such as diarrhea.^19^ However, HFL also cause unspecific toxicity in hosts, with even as little as a double dose being enough to be lethal.^20-22^ Furthermore, HFL has also been shown to spread into tissues such as muscle, organs and fat, this delays slaughter of animals, further exacerbating the financial burden on farmers.^22^ Hence, HFL does not provide a complete cure for cryptosporidiosis in claves, hence alternatives needs to be examined and developed further.^21^

It is thus paramount that more efficient treatment options to manage *Cryptosporidium* infections are explored. Majuanatha *et al*., have recently shown that a cryptosporidial lipid kinase inhibitor, KDU731, is able to efficiently kill *C. parvum* both *in vitro* and *in vivo* systems with minimal toxicity.^23^ KDU731 is a pyrazolopyridine analogue that works by competing with ATP molecules by strongly binding with the lipid kinase Phosphatidylinositol 4-kinase (PI4K) binding site of *C. parvum* parasites.^23^ More interestingly, KDU731 has been shown to eradicate cryptosporidiosis in an immunocompromised mouse model, giving a new hope for cryptosporidiosis treatment among the immunosuppressed population.^23^

In this study, we directly compared the efficacy of nitazoxanide, HFL, KDU731, and paromomycin (as a control) to the asexual growth stages of *C. parvum* using HCT-8 cells model as host cells. In addition, the toxicity and host-cell life cycle induced by the selected compounds are explored. Our aim was to bring a step closer towards understanding the efficacy of the current clinical anti-cryptosporidial compounds (and KDU731) on specific asexual stages of the parasite.

## Methods

### Parasites and host cells

*C. parvum* oocysts (IOWA strain) were obtained from the *Cryptosporidium* production laboratory (University of Arizona, USA) and stored at 4°C in Penicillin and Streptomycin (100 U/ml and 100 μg/ml, respectively; ThermoFisher Scientific #15140122) in Phosphate-Buffered Saline, pH 7.2 (PBS; ThermoFisher Scientific #21600010), for up to four months. The host cell line HCT-8, derived from human ileocecal adenocarcinoma (ATCC # CCL-225), was a generous gift from Professor Parry Guilford (University of Otago, New Zealand). HCT-8 cells were cultured in maintenance medium RPMI 1640 with GlutaMAX™ and HEPES supplements (ThermoFisher Scientific # 72400047) plus 5% foetal calf serum (FCS; CAT##) and incubated at 37°C + 5% CO_2_. When cells reached 80-90% confluency they were passaged using 0.25% trypsin/EDTA (Gibco, Cat. #25200056).

### Compounds

Nitazoxanide (Sapphire Bioscience #S1627), HFL (Wuhan Golden Wing Industry & Trade Co., Ltd), KDU731 (Novartis, Singapore), paromomycin (Sapphire Bioscience #23634) were all reconstituted in 100% dimethyl sulfoxide (DMSO; ThermoFisher Scientific #4121). Reconstituted compounds were stored at -20°C for up to 3 months before use.

### *48-hour growth inhibition assay for C. parvum in vitro* IC value determination

HCT-8 cells were seeded in either 96-well plates or 24-well plates at an inoculum of 2 × 10^4^ or 1 × 10^5^ of cells / well, respectively. HCT-8 cells were then incubated until confluency reached approximately 80% before infection. *C. parvum* oocysts were first treated with diluted household bleach (at 1:4 in water) for 10 minutes on ice to sterilise any potential contaminants before being washed with sterile Milli-Q^®^ water and centrifuged at 16,000 ×g for 3 minutes twice. The final pellet was then resuspended in 100 µL of 0.75% sodium taurocholate (Sapphire Bioscience #16215) and incubated 45 minutes at 37°C + 5% CO_2_ to allow excystation of the sporozoites. The medium of the host cells was replaced with fresh medium but with 3% horse serum (R3), and each plate wells supplemented with either 1 × 10^4^ or 1 × 10^5^ sporozoites/well for the 96-well or 24-well plates. The plates were finally spun at 150 × *g* for 3 minutes with low deceleration. Infected cells were then incubated at 37°C + 5% CO_2_. After 3 hours, all wells were gently washed with warm PBS twice and medium was resupplied to each well before incubation is resumed. Where stated, the resupplied medium was supplemented with either nitazoxanide, HFL, KDU731, or paromomycin at specified concentrations.

At specified timepoints, infected HCT-8 cells in 96-well plates were fixed with sterile 4% paraformaldehyde (PFA; VWR Chemicals #28794.295) in PBS for 10 minutes at room temperature, permeabilised with 0.25% Triton X-100 (Merk #SLBP6453V) in PBS for 10 minutes at 37°C, and then blocked with 1% bovine serum albumin (BSA; Roche #3853143) in PBS for 1 hour at room temperature. The parasitophorous vacuoles (PV) of the parasites of were then stained with 2 μg/mL fluorescein labelled Vicia Villosa lectin (VVL; Vector Laboratories #ZG0124) in 1% BSA/PBS for 1 hour at room temperature then nuclei stained with 2 μg/mL DAPI (Merk #096M-4014V) in Milli-Q^®^ water for 15 minutes in the dark. Following each stain, the wells were washed with 0.1% Tween-20 (Merk #SLBZ8563) in PBS thrice. Each well was then imaged using the Evos FL Auto 2 cell imaging system microscope (ThermoFisher Scientific) and 25% field view from the centre of each well was captured using a 20× magnification lens.

For infected HCT-8 cells grown in 24-well plates, the infected cells were first lifted using 0.25% trypsin/EDTA, centrifuged at 500 ×g for 4 minutes, and pellet resuspended in 200 μL PBS before DNA was extracted using the QIAmp DNA Mini Blood Kit (Qiagen #163043063) as per the manufacturer’s protocol and DNA was eluted in 100 μL. The DNA was then subjected to quantitative PCR (qPCR) using PrimeTime Gene Expression Master Mix (Integrated DNA Technology; IDT #1055772), primers for the *C. parvum hsp70* gene at 750 nM each (forward: 5’-AACTTTAGCTCCAGTTGAGAAAGTACTC-3’; reverse: 5’-CATGGCTCTTTACCGTTAAAGAATTCC-3’; IDT), and *hsp70* probe (5’-AATACGTGT/ZEN/AGAACCACCAACCAATACAACATC –3’; dye/que 6-FAM/ZEN/3’ IBFQ Probe; IDT) at 200 nM. Triplicate samples (at 10 µL) in MicroAmp Fast Optical 96 well reaction plates (Applied Biosystems #4346906) were then subjected to qPCR using the Applied Biosystems ViiA 7 (ThermoFisher Scientific) for 40 cycles using the following thermocycling parameters: initial polymerase activation at 95 °C for 15 min; denaturation at 95 °C for 15 seconds followed by annealing/extension at 60 °C for 1 min. To quantify parasitic growth, standards of known *C. parvum* sporozoite number (4 × 10^6^ – 4 × 10^2^) were incorporated into the qPCR plates and used to extrapolate the number of the *hsp70* gene copies from detected cycle threshold values.

### Early PV formation / trophozoite inhibition assay

HCT-8 cells grown to ≥ 90% were infected and treated simultaneously with *C. parvum* sporozoites (2 × 10^4^) and compounds at IC_90,_ respectively. A negative invasion control was performed by pre-fixing HCT-8 cells with 4% PFA for 10 minutes at room temperature. Infected cells were incubated at 37 °C with 5% CO_2_ for 3 hours to allow early PV/trophozoite formation. HCT-8 cells were then washed, stained, and imaged using the same protocol as in the standard 48-hour growth assay (see above).

### Merozoite egress determination and inhibition assay

This method was adapted from Jumani *et al*.,^24^ briefly, HCT-8 cells grown to ≥ 90% were infected with *C. parvum* as described above. Cells were then infected with 2 × 10^4^ oocysts, post excystation. To determine when the highest egress of merozoites occur, twelve timepoints were taken for the first 24 hours of infection and processed for detection via fluorescent microscopy (see above). Once the merozoite egress time has been determined, drugs, at IC_90_, were added at 3 hours post-infection (p.i.) and parasitic growth was measured at peak egress timepoint.

### Cytotoxicity assay

HCT-8 cells were seeded in a 96-well plate and incubated at 37°C + 5% CO_2_ until confluency reached 90%. Drugs were added to wells containing and plate was incubated at 37°C + 5% CO_2_ for 47 hours. Next, resazurin (Biotium #30025-2) was added as a 10% volume of sample wells or to 100 μL of media only (as a fluorescent background control). Plate was then incubated at 37°C + 5% CO_2_ for 2 hours before 60% of the wells’ supernatant was carefully transferred into a black-well plate (Greiner, #655076). A VarioskanTM LUX plate reader (Thermo Fisher Scientific #VL0000D0) was used to measure fluorescence at excitation/emission of 540nm/585nm.

### Cell cycle assay

HCT-8 cells grown in 24-well plates, until >95 % confluent in R10 media, were treated with IC90 values of compounds in R3 for 24 hours at 37 °C with 5% CO_2_. At 24 hours, samples were refreshed with R3 (no phenol red) containing compounds to mimic clinical dosing; dead cells from the supernatant were kept with centrifugation (500 × ***g*** for 4 minutes) and replaced into the culture during refreshment of drugs. At 48 hours, the cells were washed 2× with PBS, where supernatants from culture and washes were kept in 15 mL falcon tubes. Cells were uplifted gently using Accutase® (ThermoFisher, Cat. #A1110501) at RT for 10 minutes, to be transferred to the falcon tubes. Cells and their supernatants were spun at 200 × ***g*** for 4 minutes at 4 °C, followed by 1× cold PBS wash. After another centrifugation, the supernatant was removed and replaced with 1 mL of hypotonic fluorochrome solution consisting of 1% *w/v* sodium citrate, 0.1 % Triton X-100, 50 μg/mL propidium iodide (ThermoFisher Scientific, #P1304MP) and 5 μg/mL of RNase. Samples were stored in the dark at 4 °C and were analysed within one week using the BD LSRFortessa Cell Analyser using the YG_G10/20 channel (50,000 events captured). Flow cytometry data were exported and analysed using FlowJo v10: after gating for singlet populations, the biology application – Cell Cycle function, was used to obtain the percentage of cells at the G1 phase, S phase, and G2/M phase of the cell cycle.

### Statistical analyses

Statistical analysis of experimental results in this project were executed using GraphPad Prism9. All comarisons were made using one-way ANOVA with Šídák’s multiple comparisons test post corrections.

### Image J Software Code to detect PVs

Note: *set scale prior using known distances in µm to pixel values*

Run Macro using the following code:

run(“Subtract Background…”, “rolling=10”);

run(“8-bit”);

setAutoThreshold(“Huang dark”);

//run(“Threshold…”); setThreshold(5, 255);

setThreshold(5, 255);

//setThreshold(5, 255);

setOption(“BlackBackground”, false);

run(“Convert to Mask”); run(“Watershed”);

run(“Fill Holes”);

run(“Analyze Particles…”, “size=2.5-40 summarize”)

## Results

This study was set out to directly compare the *in vitro* efficacy of clinical anti-cryptosporidial compounds Nitazoxanide and HFL with the lipid kinase inhibitor KDU731 using paromomycin as a control. The asexual growth of *C. parvum* was first verified by culturing the parasites *in vitro* on HCT-8 host cells. Fluorescent microscopy or qPCR were employed to detect parasitic growth (Figure 1). For both methods and compared to the initial (3-hour p.i.), the highest parasitic growth was detected at 48 hours p.i. with fold increase of 6.3 (mean PV detected: 1613 ± 146.9 SD; Figure 1B) and 22089 (mean DNA copy number 94099 ±953.9 SD; Figure 1C and D).

**Figure 1.**
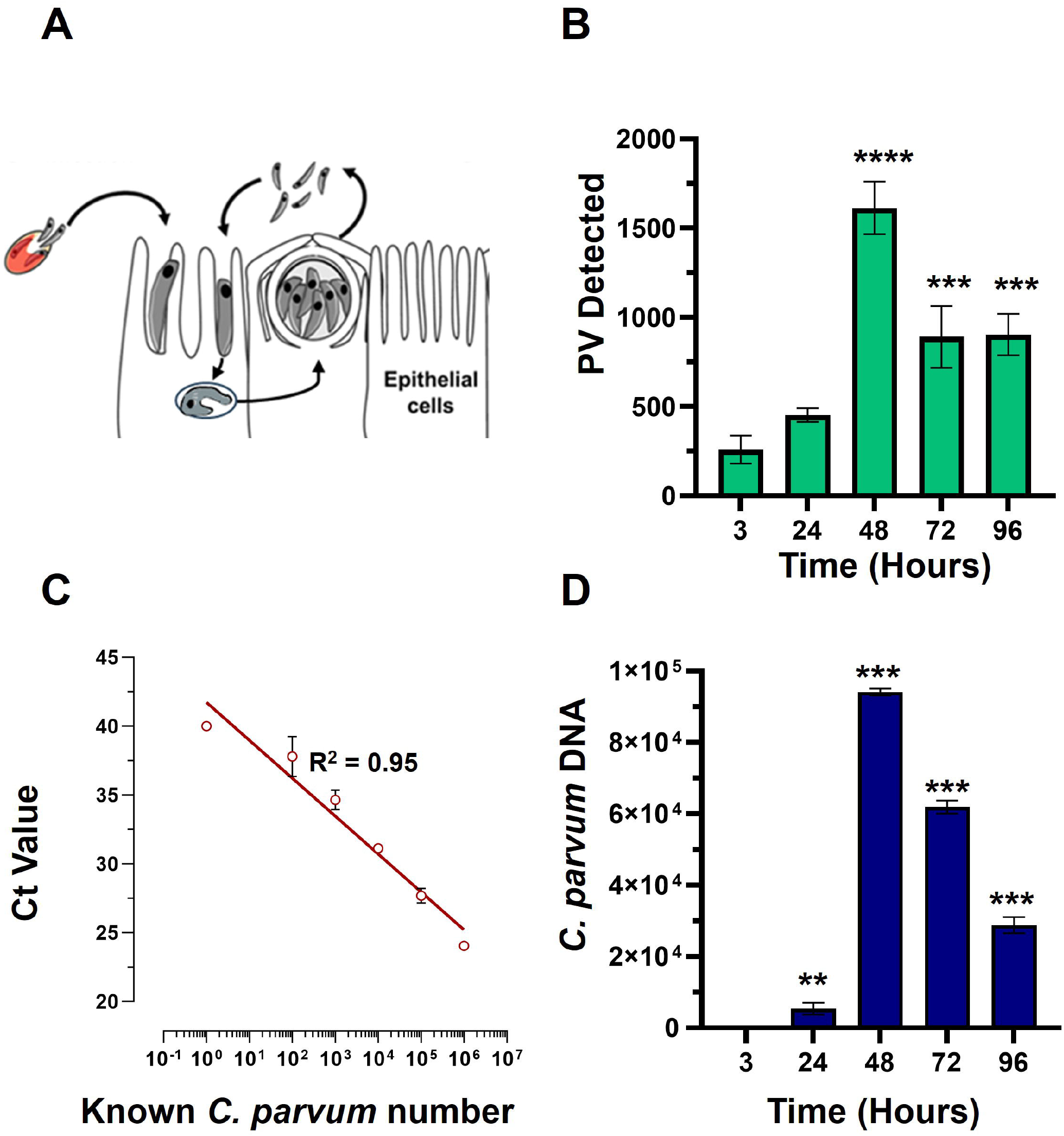
*In vitro* growth of *C. parvum* in HCT-8 cells. **(A)**HCT-8 host were infected with *C. parvum* sporozoites after excystation and growth was analysed at 3-, 24-, 48-, 72-, and 96-hours post infection (p.i.). Samples were either subjected to detection via fluorescent microscopy or qPCR. **(B)** *C. parvum* parasitophorous vacuoles (PV) detected using VVL-FITC and quantified using ImageJ. (**C & D**) *C. parvum* infected HCT-8 cells were instead lifted, lysed, and total genomic DNA extracted at given timepoints. The cryptosporidial *hsp70* gene (heat shock protein 70) was targeted using primers and a specific TaqMan probe. **(C)** Representative best fit graph showing known parasite numbers against delta CT (cycle threshold) values; R^2^ = 0.9526. (**D**) Unknown numbers of parasites were interpolated using the best fit line and CT values. Scale bar: 100 µm. Statistical analysis comparing each time point to the 3-hour time point was done using a one-way ANOVA with Šídák’s multiple comparisons test post corrections. **: *p* < 0.008, ***: *p* < 0.001.

Nitazoxanide, Paromomycin, HFL, and KDU731 were all then tested against the total growth of *C. parvum in vitro* using the 48-hour time-point to calculate the half-maximal inhibitory concentrations (IC_50_) and IC_90_. All four compounds demonstrated varying inhibition activities against *C. parvum in vitro*, as verified using both methods of detection. KDU731 displayed the lowest IC values followed by HFL, then Nitazoxanide, and finally with highest IC values shown by Paromomycin (Figure 2C). Overall, the IC_50_ values did not vary significantly between the samples detected using either microscopy or qPCR, safe for nitazoxanide which displayed a mean difference of 5563 nM between the two methods (p = 0.003; Figure 2C). The IC values for each compound as analysed using microscopy and qPCR are detailed in Table 1. Since there are lower variances (SD) between values detected using qPCR, and the method quantifies individual parasites rather than non-specific PVs, the IC values from the qPCR batch were used for the proceeding experiments. The microscopy detection method was however, chosen for the proceeding experiments as it is significantly less laborious and cost-effective.

**Table.1.** Summary of the anti-cryptosporidial efficacy of the compounds tested.

|  | Fluorescence Microscopy (PV) |  |  |  | qPCR (*hsp70*) |  |  |  |
| --- | --- | --- | --- | --- | --- | --- | --- | --- |
| Drug | IC<sub>50</sub> (nM) | SD | IC<sub>90</sub> (nM) | SD | IC<sub>50</sub> (nM) | SD | IC<sub>90</sub> (nM) | SD |
| PMC | 27720 | 2.40 | 105870 | 10.92 | 25020 | 1.29 | 52250 | 4.28 |
| NTZ | 3607 | 3.98 | 51420 | 15.81 | 9170 | 1.00 | 19860 | 3.22 |
| HFL | 117 | 6.34 | 195 | 15.03 | 77 | 3.19 | 178 | 8.39 |
| KDU731 | 91 | 2.60 | 161 | 4.99 | 102 | 2.28 | 146 | 2.92 |

*PMC: paromomycin; NTZ: nitazoxanide; HFL: halofuginone lactate* *PV: Parasitophorous Vacuole; hsp70: Heat-Shock protein-70; n=3*

**Figure 2.**
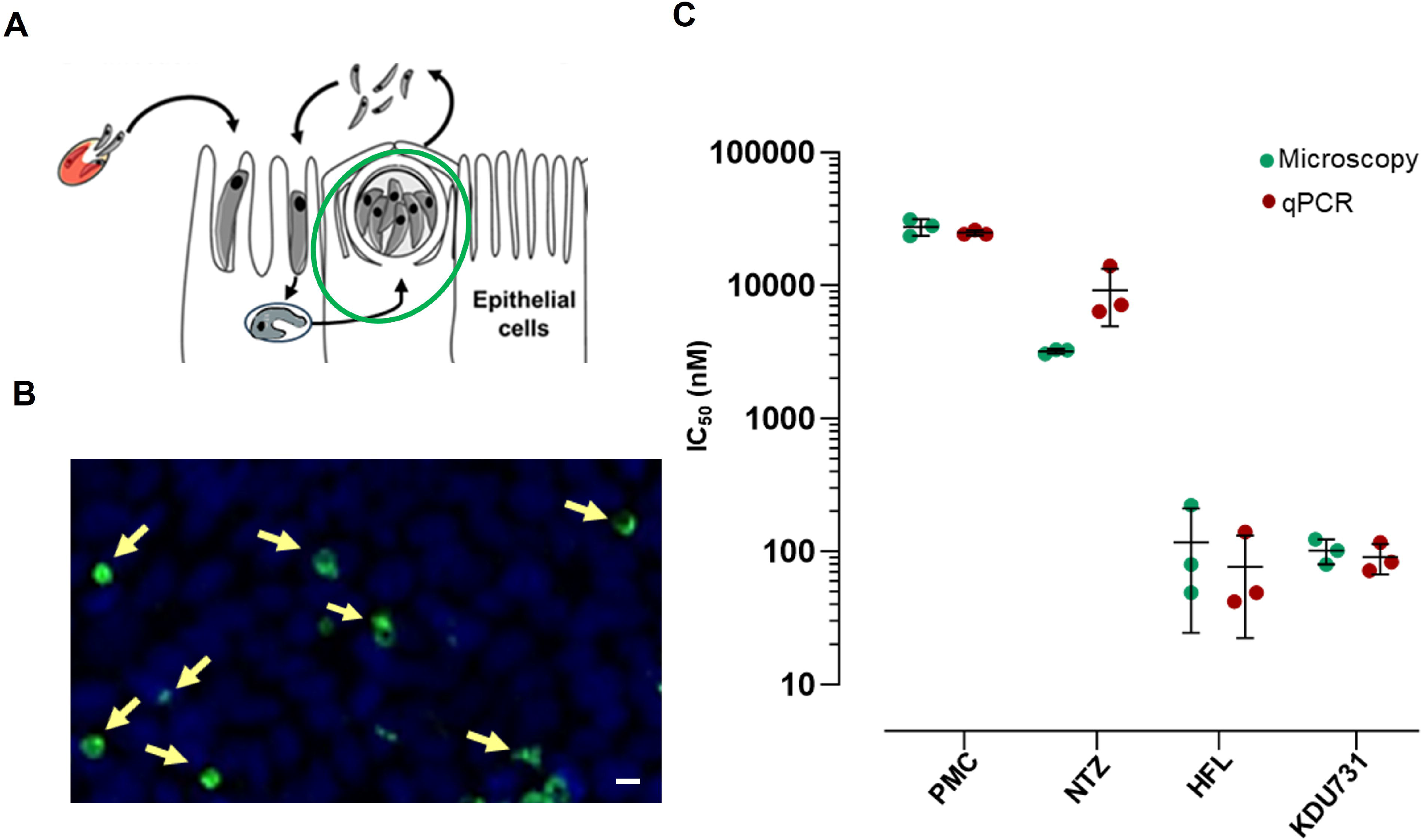
Inhibition of *C. parvum* growth by clinical compounds and KDU731 *in vitro* to estimate IC_50_. **(A-B)** Graphical and representative image of stage and PV measured 48-hours p.i. **(C)** IC_50_ values of parasites treated with each drug were determined either by measuring PV numbers of DNA copy number (microscopy or qPCR, respectively) and calculated by non-linear regression using four-parameter logistic curve. The IC_50_ values from each compound are summarised and compared between the two methods. Data representatives of three independent experiments.

Trophozoites are typically formed 3 to 4 hours p.i. (Figure 3A-B), The PV formed between sporozoite invasion and trophozoite formation is referred to here as early PV formation. To determine of any of the test compounds had an effect against early PV formation, HCT-8 cells were treated with either compound (at IC_90_; Table 1). As expected, the control with PFA-fixed cells was significantly hindered for early PV formation when compared to the no drug control (4.3% PV detected, ±4.96 SD). This was followed by nitazoxanide which also significantly suppressed early PV formation (Figure 3C; mean growth 84.11% ± 6.0 SD). No statistical significance was observed among the other compounds. The next asexual phase of *C. parvum* is the formation and release of merozoites from type I meronts. We have cultured *C. parvum* and observed the peaks of parasitic yields within a 24-hour period (Figure 4). Using qPCR, the analysis showed that the most observable peaks were at 12-, 17.5-, and 18.5-hours p.i. With the 18.5 hours p.i. yielding the highest parasitic yield (mean growth 145.4% ±1.67 SD; Figure 4B). This was next applied using the compounds we are testing, and merozoite burst/egress has been studied (Figure 4C). With an untreated mean growth ratio of 1.37, the merozoite egress was significantly suppressed in parasites treated with halofuginone lactate and PMC followed by nitazoxanide (mean growth ratio 1.03 ± 0.01 SD, 1.08 ± 0.08 SD, and 1.193 ± 0.07 SD, *respectively*). There was no statistical significance with parasites treated with KDU731.

**Figure 3.**
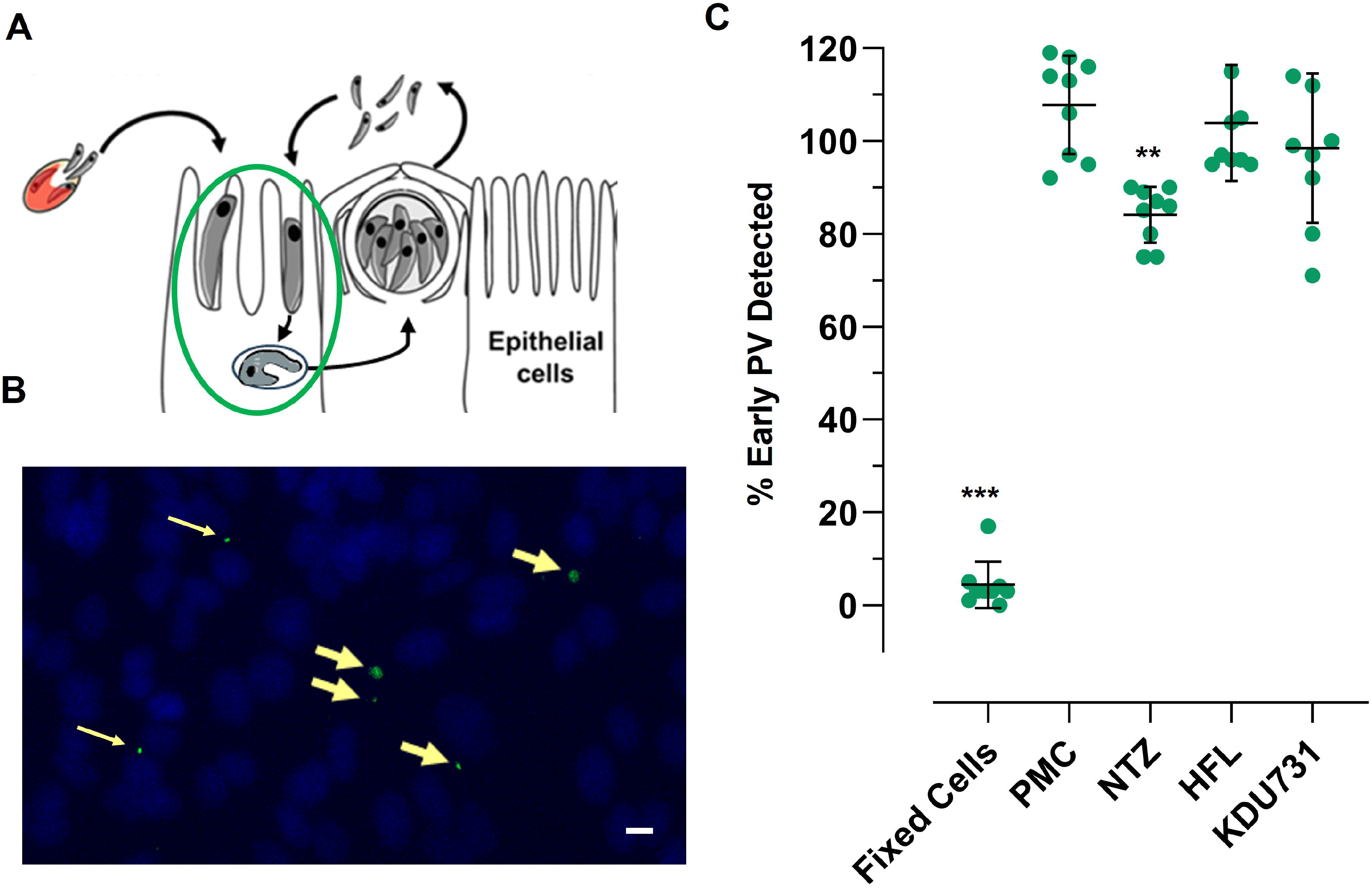
Effects of selected compounds on early PV (trophozoite) formation. Live or fixed HCT-8 cells were infected with *C. parvum* and treated with each compound at IC_90_ simultaneously. Cells were fixed with 4% PFA at 3 hours p.i. and stained with VVL-FITC (green) and Hoechst 33342 (blue). Cells were analysed using fluorescent microscopy and PVs were counted using ImageJ software. **(A-B)** Graphical and representative image showing early PV formation (yellow arrows). Scale bar = 32 µm **(C)** Quantified PV were plotted and analysed using one-way ANOVA with Šídák’s multiple comparisons test post corrections, comparing each to the control (DMSO only) sample. **: *p*= 0.007; ***: *p* <0.001. Representative of three independent experiments.

**Figure 4.**
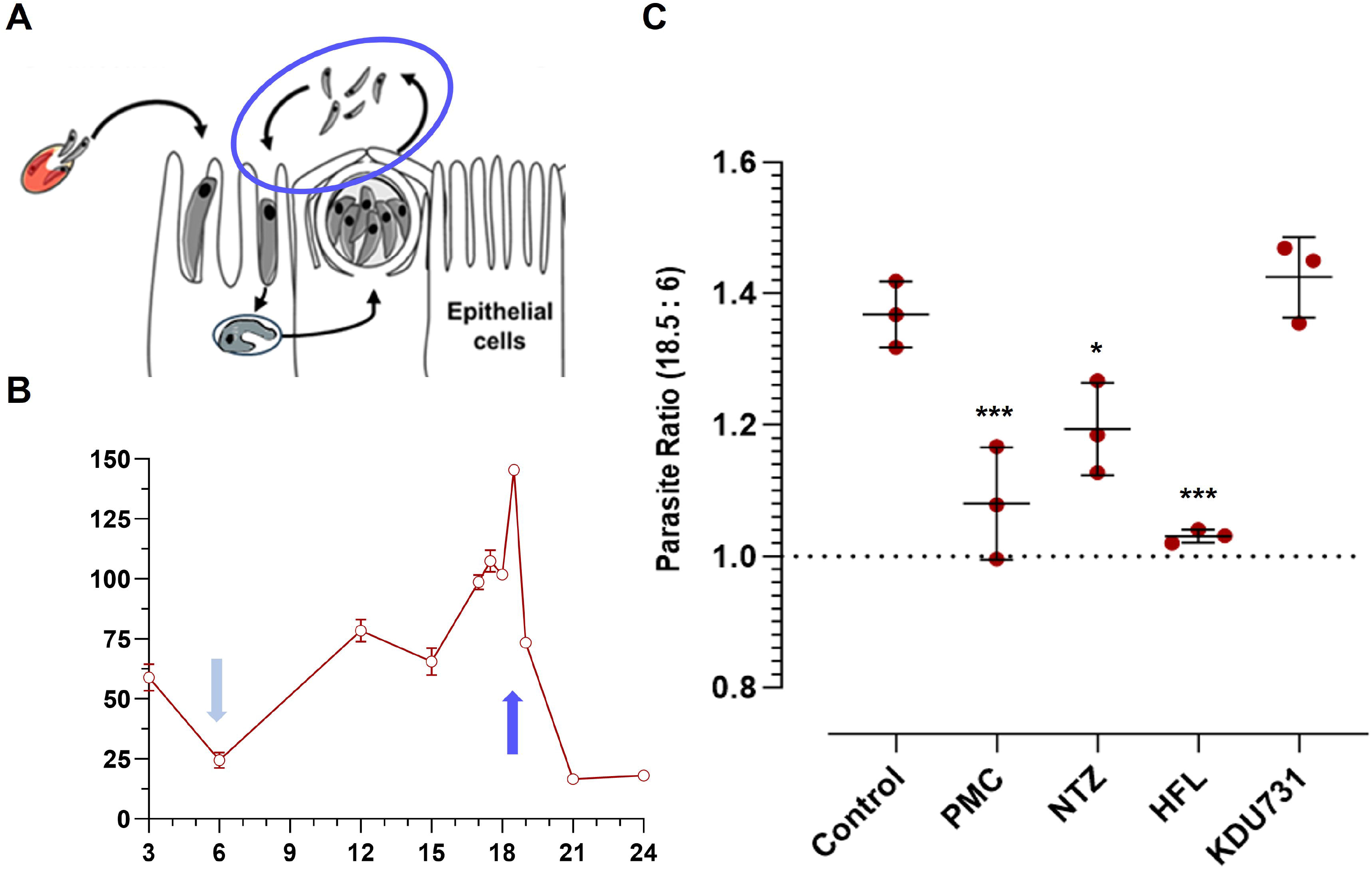
Comparison of compounds’ effects on *C. parvum* merozoite egress. **(A-B)** HCT-8 cells were infected with *C. parvum* sporozoites and parasites enumerated using qPCR at specified timepoints. Three peaks were identified at 12-, 17.5-, and 18.5-hours p.i. and compared to the lowest yielding timepoint (6-hour). **(C)** Ratio of parasites detected between 18.5-and 6-hours p.i. samples treated with either of the compounds at IC_90_. All data was analysed using one-way ANOVA with Šídák’s multiple comparisons test post corrections, comparing each to the control (DMSO only) sample. *: *p* = 0.02; ***: *p* < 0.001. Results are representative of two independent experiments.

To determine if the compounds affect the viability of host cells, the metabolic capacity and growth cell cycle of the host cells were examined. The metabolic capacity was analysed using a resazurin assay using two treatment regiments: a one-off dose or a continuous daily dose of each compound, all at IC_90_ (Figure 5A). In both regiments, PMC displayed the lowest cell viability (mean growth one dose: 38.6% ±8.17 SD; continuous dose: 24.87% ± 20.18 SD). Due to the high variability, nitazoxanide displayed a modest decrease in host cell viability (mean growth 74.7% ±24.16 SD) but did not reach statistical significance. There was no significant viability change in host cells treated with either halofuginone lactate or KDU731. To verify the safety of halofuginone lactate and KDU731, the host cells were treated with either drug at IC_99_ for up to 72 hours (Figure 5B). There was only a significant drop in host cell viability when treated with halofuginone lactate compared to the control (p < 0.001).

**Figure 5.**
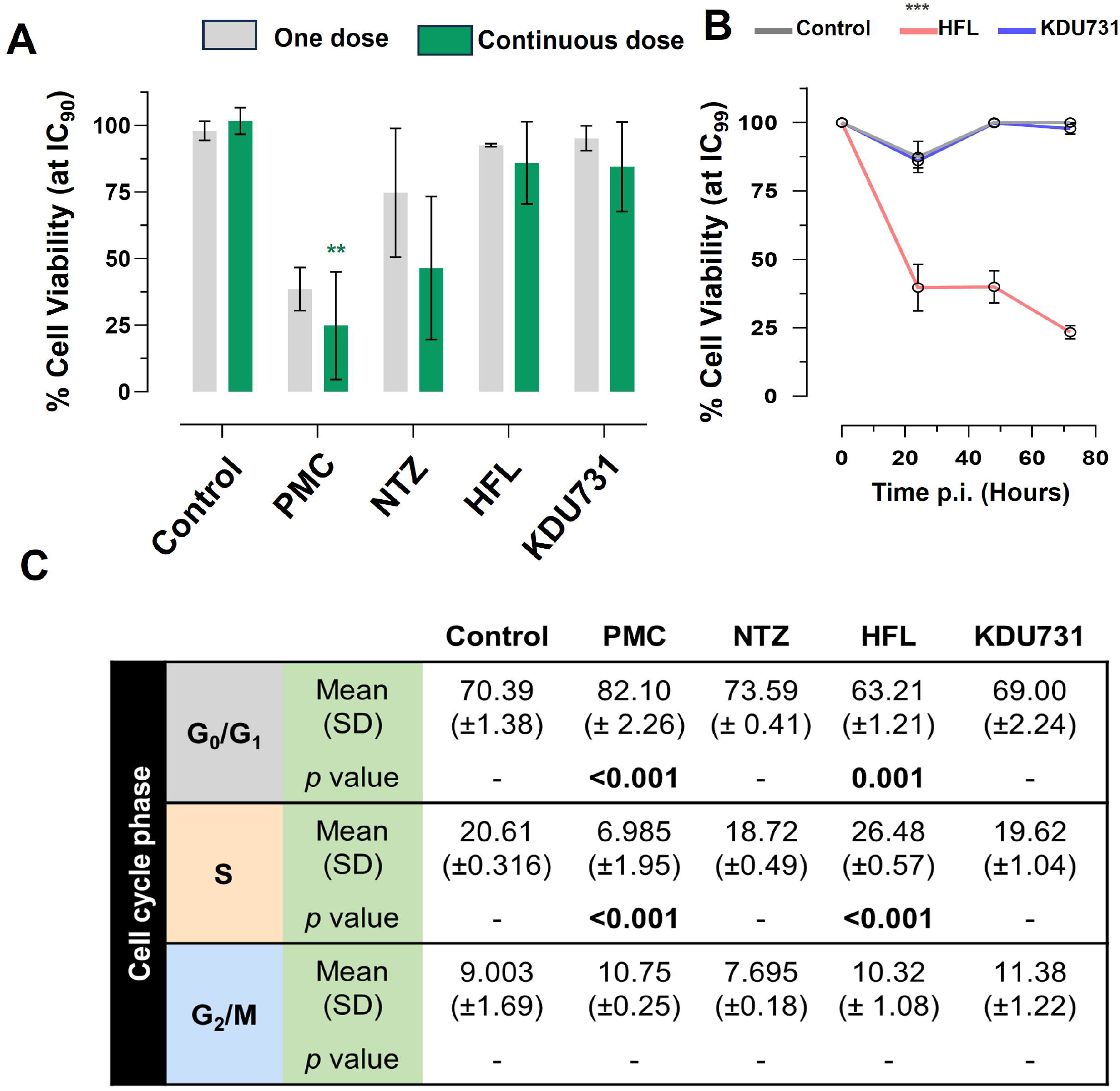
Determining toxicity of the selected compounds on HCT-8 host cells using Resazurin assay and cell cycle analysis. **(A)** HCT-8 cells (at 80-90% confluency) were treated with each compound at IC_90_ and incubated at 37°C + 5% CO_2_ for 48 hours. Cells were either treated with the compounds once (day zero; one dose), or daily in fresh medium (continuous dose). At 47 hours post incubation, resazurin was added to the wells and incubated at 37°C for 1 hour and fluorescence was measured (excitation/emission 540/585 nm) **(B)** Cells were instead incubated with either HFL or KDU731 at IC_99_ (or DMSO only control) for up to 72 hours and viability measured via Resazurin. The area under the curve for each sample was calculated and plotted. **(C)** Alternatively, after the incubation with all compounds at IC_90_, cells were then lifted and strained with hypotonic fluorochrome solution containing propidium iodide. The cells were then subjected to flow cytometry. Singlet populations were gated, and fluorescence detected using the YG_G10/20 filter channel (50,000 events captured). Flow cytometry data were analysed using the FlowJo v10 software: using the Cell Cycle function tool to obtain the percentage of cells undergoing growth (G1), DNA synthesis (S), and pre-mitosis growth phase (G2). Mean values from each cycle phase were compared using one-way ANOVA with Šídák’s multiple comparisons test post corrections, comparing each to the control cell population (DMSO only). Results are representative of two independent experiments. PMC: paromomycin; NTZ: nitazoxanide; HFL: halofuginone lactate.All statistical analyses were done using one-way ANOVA with Šídák’s multiple comparisons test post corrections, comparing each to the control (DMSO only) sample. *: *p* = 0.02; **: *p* = 0.002; ***: *p* < 0.001. Results are representative of three independent experiments.

The effects of each drug on the growth cell cycle of the host cells were then investigated (Figure 5C). The drug concentrations selected were from the IC_90_ because cells subjected to IC_99_ concentration displayed very high toxicity for halofuginone lactate ad there were low number of cells to analyse. There was a significant increase in cells undergoing the G_1_ phase when treated with PMC (mean difference 11.71%) and a significant decrease at the S phase compared to the control (mean difference 13.62% ± 1.95 SD). In contrast, halofuginone lactate significantly decreased cells undergoing G_1_ phase (mean difference 7.18%) and a significant increase of cells undergoing the S phase compared to the control (mean difference 5.87%). No statistically significant change was observed among cells treated with nitazoxanide or KDU731 (Figure 5C).

## Discussion

The findings of this study shed light on the varying efficacies and safety profiles of Nitazoxanide, HFL, KDU731, and Paromomycin against *C. parvum*. Notably, KDU731 displayed the most promising profile, with low nanomolar activity and low host cell toxicity. This aligns with its mechanism of specifically targeting *Cryptosporidium* lipid kinase PI(4)K (phosphatidylinositol-4-OH kinase), which seems to have minimal impact on merozoite egress.^23^ In contrast, Nitazoxanide and HFL, while effective in certain stages, showed variability in results and a degree of toxicity to host cells.

An important distinction was observed in the methodology of parasite detection. Microscopy, measuring PVs, and qPCR, quantifying individual parasites, provided different insights. Particularly, Nitazoxanide displayed notable variability between these methods, suggesting potential inconsistencies in its measurement.

The study’s findings on early PV formation and merozoite egress contribute valuable information to the understanding of the compounds’ stage-specific actions. Notably, Nitazoxanide significantly suppressed early PV formation, indicating its potential in the early stages of infection. However, its high variability and modest impact on host cell viability call for caution.

In terms of host cell viability and safety, KDU731 stands out for its minimal toxicity at both IC_90_ and IC_99_ concentrations, unlike Paromomycin and Nitazoxanide. This aspect is crucial, as the safety of the host cells is a key concern in the treatment of cryptosporidiosis. Furthermore, the study reveals that HFL can alter the host cell cycle, thereby highlighting the superior profile of lipid kinase inhibitors like KDU731 in terms of safety and minimal impact on host cells.

The clinical drugs Nitazoxanide, paromomycin, and halofuginone lactate can also have a negative impact on patients’ intestinal microflora. For example, paromomycin is known to cause significant changes in the gut microflora of patients after treatment.^25^ Moreover, a recent study showed that *C. parvum* infections significantly decreases the alpha diversity of the gut microbiome in humans and mice.^26,27^ It is therefore important to consider not only the anti-cryptosporidial effects of the compounds but also the conservation of an optimal gut microbiome profile. The effects of KDU731 on the gut microbiome is yet to be elucidated, but since it is not a broad-range antimicrobial such as paromomycin or Nitazoxanide, then it is expected to have minimal microbiome effects.

Overall, this study emphasizes the need for more efficient and safe treatment options for cryptosporidiosis. The promising profile of KDU731, particularly in terms of efficacy and safety, positions it as a potential leading candidate in the fight against Cryptosporidium infections. The findings underscore the importance of considering both the efficacy against the parasite and the safety for the host in developing anti-cryptosporidial therapies.

